# Tensorial Independent Component Analysis Reveals Social and Reward Networks Associated with Major Depressive Disorder

**DOI:** 10.1101/2022.08.04.502820

**Authors:** Jeff B. Dennison, Lindsey J. Tepfer, David V. Smith

## Abstract

Major depressive disorder (MDD) has been associated with changes in functional brain connectivity. Yet, typical analyses of functional connectivity, such as spatial ICA for resting-state data, often ignore sources of between-subject variability, which may be crucial for identifying functional connectivity patterns associated with MDD. Typically, methods like spatial ICA will identify a single component to represent a network like the default mode network (DMN), even if groups within the data show differential DMN coactivation. To address this gap, this project applies a tensorial extension of ICA (tensorial ICA)—which explicitly incorporates between-subject variability—to identify functionally connected networks using fMRI data from the Human Connectome Project (HCP). Data from the HCP included individuals with a diagnosis of MDD, a family history of MDD, and healthy controls performing a gambling and social cognition task. Based on evidence associating MDD with blunted neural activation to rewards and social stimuli, we predicted that tensorial ICA would identify networks associated with reduced spatio-temporal coherence and blunted social and reward-based network activity in MDD. Across both tasks, tensorial ICA identified three networks showing decreased coherence in MDD. All three networks included ventromedial prefrontal cortex (vmPFC), striatum, and cerebellum and showed different activation across the conditions of their respective tasks. However, MDD was only associated with differences in task-based activation in one network from the social task. Additionally, these results suggest that tensorial ICA could be a valuable tool for understanding clinical differences in relation to network activation and connectivity.

## Introduction

Major depressive disorder (MDD) is a psychiatric disorder that has been associated with disruptions to reward processing (Cooper et al., 2018) and social cognition (Weightman et al., 2019). These differences in processing rewards and social stimuli have been linked to changes in the activation of specific brain regions using functional MRI (fMRI). For example, compared to healthy controls, individuals with MDD have greater reward-dependent activation of the ventromedial prefrontal cortex (vmPFC) (Ng et al., 2019) and middle frontal gyrus (MFG) (Zhang et al., 2013) but less activation of the striatum (Ng et al., 2019). However, in response a social stimulus, the opposite pattern has emerged: perception of social stimuli in MDD is associated with greater activation of the striatum and less activation in vmPFC (Lai, 2014). The disruptions of activation within frontal cortex and striatum across reward and social tasks suggest that MDD could also be associated with the disruption of both reward- and social-evoked connectivity between striatum and vmPFC.

Building on these observations of aberrant patterns of brain activation, other studies have shown how MDD is associated with differences in functional connectivity. Additionally, differences in task-dependent connectivity associated with MDD using both rewards and social manipulations have converged on a common set of regions. Task-dependent connectivity between the striatum and ACC has been shown to be relatively less for individuals with MDD in response to rewards (Morgan et al., 2016; Admon et al., 2015) and greater in response to social stimuli (Healy et al., 2014). Connectivity has also been shown to be altered between the striatum and vmPFC under depression in response to both rewards (Furman et al., 2011;Hanson et al., 2018) and social stimuli (Tepfer et al., 2021). Though these results suggest that connectivity from the striatum to both the ACC and vmPFC differ for those with MDD, this could either imply that MDD is associated with changes to a single larger network which incorporates each of these areas or two independent networks, one highlighting activity in the striatum and vmPFC and the other striatum and ACC.

While it can be difficult to resolve how many networks are involved in a behavior using seed-based approaches (Smith et al., 2014), Independent Components Analysis (ICA) has been used to identify independent but partially-overlapping networks consistent with theory as we do not expect brain networks to contain mutually exclusive regions (Smith et al., 2012). For example, if the striatum was coactive with both the mPFC and amygdala but the mPFC and amygdala were not coactive with each other, ICA could then identify two independent networks (i.e., striatum-mPFC and striatum-amygdala networks). Indeed, early research in a simulated driving task identified two networks which overlapped in the mPFC, but only one had increased activation during the driving portion of the task (Calhoun et al., 2002).

Rather than predefining a seed a priori, which can bias what regions are shown to have functional connectivity(Cole et al., 2010), ICA is a data driven approach that has more relaxed assumptions. ICA identifies different networks by creating a set of linear combination of voxels (i.e., networks) which has the greatest independence in the resulting time courses (see Calhoun et al., 2010 for a description of the different algorithms that can be used). This approach can be especially advantageous when a researcher does not want to define a seed region or have a pre-defined time-course for a network.

Though ICA based methods have some advantages over seed-based approaches, investigating individual differences using ICA required additional methods to be developed, such as dual regression (Filippini et al., 2009; Nickerson et al., 2017). The dual-regression approach takes group-level ICA maps and identifies participant-level maps via two independent regressions (i.e., spatial regression followed by a temporal regression).”. However, the initial group ICA may fail to separate signals from different sub-groups in the sample as participant’s data is often concatenated either spatially or temporally removing the inter-subject information ((Beckmann and Smith 2005). For example, by not considering the group information, spatial ICA could potentially reveal a network that is present in neither healthy controls nor a clinically depressed population, but represents some mixture of the two groups. A researcher might use this network to estimate individual differences in network activity or connectivity and either fail or succeed in finding group differences despite that the initial output was a poor representation of either group in the first place.

To better separate out these group signals, we turn to a tensorial extension of ICA, which estimates the networks jointly in terms of its response across subjects, time, and voxels. This tensorial extension has been shown to reduce interference between the different estimated sources (Beckmann and Smith 2005), which may make it more suited to estimating networks with different samples (e.g., a clinical depression and healthy control group). This method produces a set of weights across voxels indicating the spatial distribution of activation, timepoints indicating the time course of the network activation, and subjects indicating each subject’s spatio-temporal coherence with the network. For example, a reward network may show the greatest weights in reward sensitive neural regions like the ventral striatum and vmPFC, the greatest temporal weights during the anticipation and consumption of rewards, and have the greatest subject weights associated with subjects who are more reactive to rewards.

While most group ICA methods only require that subject’s fMRI data is processed into a standard space, tensorial ICA also requires a common time course (i.e. the timing of events needs to be the same across subjects) precluding it from use with resting-state data. This requirement has generally limited the application of tensorial ICA, but one previous study explored networks involved in Alzheimer’s disease (Rombouts et al., 2007). This study included a sample of healthy older adults, an at risk group with mild cognitive impairments, and individuals diagnosed with Alzheimer’s disease performing a face encoding task to assess episodic memory. Applying tensorial ICA to the fMRI data identified different networks which were described by a spatial, temporal, and subject mode. By analyzing the subject mode, which represented each subjects spatio-temporal coherence with the network, for group differences Rombouts et al., was able to identify various networks which had a diminished response in dementia. Additionally, Rombout et al. showed that tensorial ICA was better able to identify group differences compared to a model-based analysis, by applying both analyses and comparing the results. Yet, this method has not been applied to psychiatric disorders such as depression.

Using tensorial ICA may also help to identify aberrant social and reward network activity in MDD. The major aim of this paper is to assess whether tensor-based ICA can help to identify differences in brain-network activation associated with MDD. To find these differences, we analyzed BOLD data from the human connectome project which gave us a large sample of participants diagnosed with MDD, healthy controls (HC), and those with a family history (FHx) of MDD. By incorporating a family history group our analyses may be able to suggest networks that represent factors involved in developing or protect against clinical depression. We chose to analyze BOLD data while participants performed a gambling and social cognition task as MDD has been associated with behavioral and neurological disturbances to reward (Ng et al., 2019; Whitton et al., 2015) and social cognition processes (Bora & Berk, 2016; Lai, 2014). Hypotheses and analyses were preregistered using the AsPredicted platform (https://aspredicted.org/BIOJHP). We predicted that networks differentiating our HC, FHx, and MDD groups across both tasks would contain the vmPFC, striatum, and cerebellum. Additionally, we predicted that group differences would be associated with differential activation to rewards and social stimuli. To compare our results, we followed our tensorial ICA analyses with two forms of dual-regression, whole brain and spatially restricted, which both rely on a traditional temporally concatenated ICA in the initial steps.

## Methods

### Participants

Due to the open-data nature of the Human Connectome Project, sample size was determined by including as many participants as possible who met the inclusion criteria for the MDD, FHx, and Healthy Control groups and used an identical sample as our previous work (see Tepfer et al., 2019 for details). Though no formal power analysis was conducted a priori, including the full set of identified participants maximized the theoretical statistical power.

Specifically, the sample included 279 participants (males, 120; females, 159; ages, 22-36; mean ± SD, 28.45 ± 3.75 years) from the 1200 Subjects Release (WU-Minn HCP consortium S1200 Data) of the Human Connectome Project (HCP). Participants were categorized into one of three groups; the MDD (N=71) group for individuals who indicated a lifetime history of major depression, the family history (FH) group (N=103) for those with a parent who had indicated a lifetime history of major depression, or the healthy control (HC) group for whose parents had not received a DSM-IV diagnosis of major depression and had indicated no family history of psychiatric or neurological disorders (N=105). See Tepfer et al. 2021 for additional details of data selection.

### Task

Participants from the HCP performed seven tasks during fMRI of which we analyzed two: the social cognition (Castelli et al., 2000) and incentive processing task (Delgado et al., 2000). Each task involved alternating blocks of stimuli. The descriptions of the two tasks of interest here are adaptations from details provided by Barch et al., 2013).

### Incentive Processing

The incentive processing task was adapted from a card guessing game task (Delgado et al., 2000). The participants guessed the number on a mystery card (represented by a “?”) in order to win or lose money. The number on the mystery card could range from 1 to 9. Participants indicated whether they thought the mystery card number would be more or less than 5 by pressing one of two buttons on the response box. Trials were pre-determined to be reward, loss, or neutral feedback trials. After making their response, participants would see the mystery number they were meant to guess along with an arrow and monetary amount to indicate whether the outcomes were rewards, losses, or neutral. On reward trials, participants would see that they correctly guessed whether the value of the card was greater or less than 5 along with a green up arrow and “$1”. Loss trials showed that the participant guessed incorrectly along a with red down arrow next to “- $0.50”. On neutral trials the mystery number was shown to be 5 and after a gray double headed arrow was displayed. Blocks of 8 trials were presented that are either mostly rewarding or mostly losses (e.g., during the reward block 6 trials of reward outcomes interleaved with either 1 neutral and 1 loss outcomes, 2 neutral outcomes, or 2 loss outcomes). See Barch et al., 2013 for details.

### Social Cognition Task

The social cognition task was adapted from a mentalizing task (Castelli et al., 2000) which involved participants presented with short video clips (20 s) of shapes (squares, circles, triangles) interacting socially or moving randomly. After each video, participants chose to indicate whether they thought the objects had a social interaction, were moving randomly, or whether the participant was not sure. Each of the two task runs has 5 video blocks (2 Mental and 3 Random in one run, 3 Mental and 2 Random in the other run) and 5 fixation blocks (15 s each). Of note, the video clips were shortened to 20 s (the Castelli et al. clips were originally 40 s) by either splitting the videos in two or truncating them. The authors deployed a pilot group to ensure that the shortened videos elicited similar responses to the full-length clips. Participants made ratings about the presence or absence of mental interactions in the videos, and these short-clip ratings were compared to the long-clip ratings for equivalence.

### Neuroimaging Data Acquisition & Preprocessing

Neuroimaging data were collected as part of the Human Connectome Project (HCP) using a 3 Tesla Siemens scanner equipped with a 32-channel RF head coil. Functional images sensitive to blood-oxygenation-level-dependent (BOLD) contrast were acquired using a Gradient-echo EPI sequence with a multiband acceleration factor of 8 [repetition time (TR): .72 s; echo time (TE): 33.1 ms; matrix 104 × 90; voxel size: 2.00 × 2.00 × 2.00 mm; 72 slices (15% gap); flip angle: 52°]. Each task was recorded in two runs with different phase encodings (either left-to-right or right-to-left). To facilitate co-registration and normalization of functional data, we also collected high-resolution T1-weighted structural scans with parallel imaging factor (iPAT) factor of 2 (TR: 2.4 s; TE: 2.14 ms; matrix 320 × 320; voxel size: 0.7 mm^3^; 256 slices; flip angle: 8°) and B0 field maps (TR: 645 ms; TE1: 4.92 ms; TE2: 7.38 ms; matrix 74 × 74; voxel size: 2.97 × 2.97 × 2.80 mm; 36 slices, with 15% gap; flip angle: 60°) as well asT2-weighted structural images with parallel imaging factor (iPAT) factor of 2 (TR: 3.2 s; TE: 565 ms; matrix 320 × 320; voxel size: 0.7 mm^3^; 192 slices; flip angle: 52°) (Barch et al., 2013).

In addition to the minimal preprocessing already featured in the HCP dataset, artifacts were identified and removed via ICA-AROMA (Pruim et al., 2015) smoothed with a 2mm fwhm kernel and runs were temporally concatenated before being submitted to tensorial ICA and dual regression. After initial analyses, the largest components contained artifacts suggesting that the network estimation was significantly influenced by large mean shifts in the signal due to the opposing phase encodings of each run (i.e., the signs of weights in the spatial mode were flipped bilaterally and a large mean shift in the time course corresponded with the end of the first and beginning of the next run). To address this issue, each run image was temporally normalized using fslmaths Tmean and Tstd options before being concatenated and submitted to further analyses.

### Neuroimaging Analyses

We used three techniques to attempt to characterize the task-dependent networks of activation and assess individual differences in connectivity. We start our analyses with tensorial ICA, which has the potential to better identify individual differences in functional connectivity than similar but more popular methods. To compare our results from tensorial ICA, we conduct two follow-up analyses relying on traditional ICA, including a whole-brain and spatially restricted dual-regression (Fig. 1).

**Figure 1:**
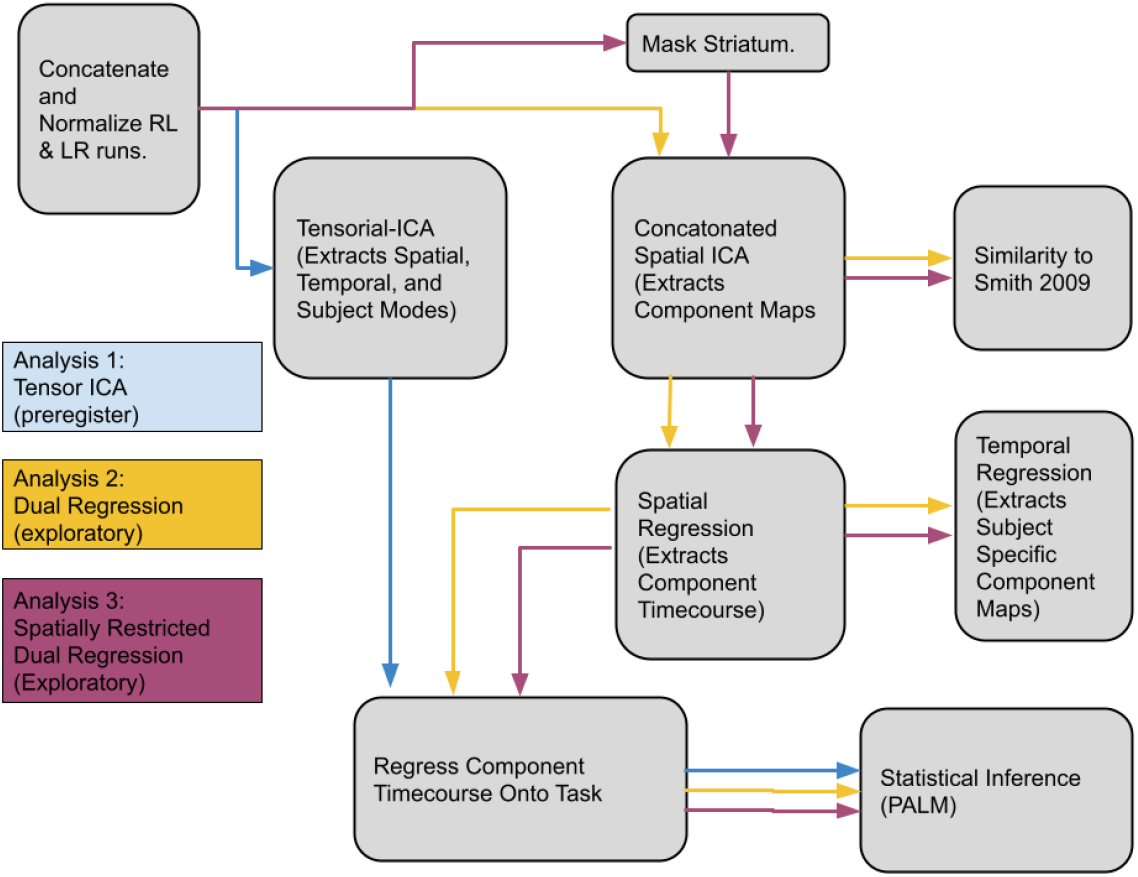
Schematic of the three analytical methods. The process for tensorial ICA (blue), whole-brain dual regression (yellow), and spatially restricted dual regression (purple) follow the arrows from each box indicating separate steps of the analysis.

### Tensorial Independent Components Analysis

Tensorial-ICA was performed using the tensorial extension through FSL’s MELODIC and set to return 25 components (Beckmann & Smith, 2005). Tensorial ICA extracts spatial, temporal, and subject modes which can be used to test our hypotheses about which brain regions, events, and participants were most highly featured in a single functional network. To identify which brain regions were involved in a network, the weights from the spatial mode were projected back into the image and a threshold was applied using a gaussian mixture model (Beckmann & Smith, 2005) to create a network map. These network maps were used to manually characterize noise and signal components using established heuristics based on the spatial features of the independent components (ICs) (Griffanti et al., 2017).

The components extracted from tensorial ICA represent networks of co-activation throughout the task. However, these networks will not all necessarily reflect co-activation in response to the different conditions in the task. It is possible that these networks reflect co-activation with aspects that are present in both conditions of a task or other features which are temporally consistent across participants (e.g., fatigue). To identify networks which are differentially activated in the different conditions of the task, the temporal modes of components identified as signals were then fit to the block design of the corresponding task (Gains vs Losses, or Mental vs Random) to gauge how each network tracked different aspects of the task. The designs were fit using an ordinary least-squares linear regression approach implemented in *statsmodels api* (https://statsmodels.sourceforge.net/) in python for the temporal modes of each participant provided by the output of FSL’s MELODIC tensorial ICA. The temporal modes were regressed onto two vectors representing the different block conditions of each task convolved with a double-gamma hemodynamic response function. The difference of the resulting betas (e.g. βGains – βLoss) were submitted to a one sample t-test using FSL’s PALM (Winkler et al., 2014) (see below for details). We then tested for group differences using a two-sample t-test with both the extracted weights from the subject modes of the tensorial ICA and also our estimates of the task-evoked responses in each subject. The subject mode reflects a subject’s overall coherence with the combined spatial and temporal features of the network. However, by testing the difference in task evoked responses we can determine if those differences are driven in part by reward or social responses. Two-sample t-tests were also conducted using PALM. For each reported effect-size, we used SciPy (Virtanen et al., 2020) to perform a reverse percentile confidence interval for 95% confidence performed using 9999 resamples (Efron & Tibshirani, 1993).

### Whole-Brain Dual Regression Analysis

Dual-regression analyses utilized a three-stage approach (see Smith et al., 2014), starting with group ICA followed by spatial-regression and temporal-regression. The data were temporally concatenated and submitted to FSL’s Multivariate Exploratory Linear Optimized Decomposition into Independent Components (MELODIC) to extract 25 independent components and returned 25 group spatial maps. During *spatial*-*regression*, spatial maps were regressed onto each participant’s functional data, resulting in a T (time points) × C (components) set of beta coefficients that characterize, in each subject, the temporal dynamics for each spatial network. Next, the data was submitted to *temporal*-*regression*, where the temporal dynamics resulting from the spatial-regression are regressed onto each subject’s functional data. This produces a set of spatial maps, specific to each subject, reflecting each voxel’s connectivity with each network identified with the group ICA. The resulting subject-specific connectivity maps reflect individual differences in connectivity such that a given network can include different regions for one individual than another and regardless of whether that brain region falls within the set of regions highlighted by the group spatial maps. Importantly, the temporal-regression step estimates each voxel’s connectivity with each spatial network while controlling for the influence of other networks—some of which may reflect artifacts, such as head motion and physiological noise.

Our core analyses were conducted on 10 well-characterized resting state networks (RSN’s) postulated to reflect cognitive and sensory functions (Smith et al., 2009). To identify RSNs from our ICA that correspond to the 10 RSNs reported in Smith et al. (2009), we conducted a spatial correlation analysis between the group spatial maps and those from Smith et al., (2009) retrieved from Oxford Centre for Functional MRI of the Brain (RRID:SCR_005283). Within both tasks, we selected the 10 components that were most highly correlated with the 10 RSNs in Smith et al. (2009) (See Table 1). The individual time-courses were regressed onto the same two vectors described for the tensorial ICA analyses representing the different conditions of each task. The difference of the resulting betas (e.g. βGains – βLoss) were submitted to a one sample t-test. For dual regression analyses, group differences were tested using a two-sample t-test using the difference between individual betas for each task resulting from the analysis of the network time course.

**Table 1:** Spatial Correlation Between Identified Components and Canonical Networks.

| Analysis | Task | Vis1 | Vis2 | Vis3 | DMN | Cer | SM | Aud | ECN | RFPN | LFPN |
| --- | --- | --- | --- | --- | --- | --- | --- | --- | --- | --- | --- |
| Whole Brain | Gamble | 0.76 | 0.6 | 0.59 | 0.66 | 0.28 | 0.38 | 0.4 | 0.42 | 0.57 | 0.65 |
|  | Social Cognition | 0.78 | 0.7 | 0.56 | 0.5 | 0.32 | 0.35 | 0.44 | 0.52 | 0.56 | 0.46 |
| Spatially Restricted | Gamble | 0.28 | 0.27 | 0.34 | 0.22 | 0.02 | 0.26 | 0.4 | 0.16 | 0.46 | 0.46 |
|  | Social Cognition | 0.36 | 0.26 | 0.41 | 0.22 | 0.05 | 0.27 | 0.39 | 0.3 | 0.42 | 0.44 |

### Spatially Restricted Dual Regression

The spatially restricted dual-regression analyses used the same three-stage approach as the whole-brain dual regression with one significant change (Leech et al., 2012). Prior to the initial group ICA the BOLD was masked so that only activation from voxels within the striatum were considered when estimating the independent components. The striatal mask was produced anatomically using the Harvard-Oxford atlas (RRID:SCR_001476) including the bilateral nucleus accumbens, caudate, and putamen.

RSN’s and task-dependency were identified using the same methods described for the whole-brain dual regression.

### Permutation-based Analysis

Output from all three analyses would reveal multiple components requiring multiple comparisons, for which we needed to control the family-wise error (FWE) or the probability of making one or more Type I errors (Voelkl, 2019)https://www.sciencedirect.com/science/article/pii/S0003347219302040?via=ihub. Additionally, several subjects within the HCP are genetically related, including some monozygotic and dizygotic twins, whose shared variance with each network may be difficult to estimate (Van Essen, et al., 2012). To control for multiple comparisons as well as the family structure, we conducted permutation-based analyses implemented with FSL’s PALM (Winkler et al., 2015). For hypotheses about the temporal mode or experimental conditions we conducted a one sample t-test on the difference between beta weights resulting from the fit of each participant’s network time-course to the task design. For hypotheses of group or individual differences we conducted two-sample t-tests submitting the difference between beta weights for each task condition. For tensorial ICA we additionally submitted individual weights from the resulting subject mode.

Code for the analyses is openly available (https://github.com/jbdenniso/CSNmaps).

### Reproducibility and Reliability

In order to assess the possible reproducibility of our results we include two additional analyses.

To better gauge the reproducibility of our findings we conducted an additional analysis on networks surviving FWE correction through PALM. For each component, we assess the ability to classify individuals into the HC or MDD group using a linear analysis. A three-fold validation procedure is used training the classifier on two thirds and determining accuracy on the remaining third of the data, taking the mean accuracy across all three folds. This is combined with a permutation test score to evaluate the significance of our cross-validated accuracy. Targets are permuted to generate ‘randomized data’ and the accuracy of a discriminant analysis is analyzed using the same three-fold validation. This is done ten thousand times to obtain an empirical null distribution of the accuracy. A p-value is then computed against the null hypothesis that features and targets are independent.

Reliability measures have been explored for typical ICA approaches. However, to better gauge the measurement reliability of our tensor ICA components, we conducted a split halves reliability test. Tensorial ICA was performed independently on both runs. After removing components containing noise-related artifacts (Griffanti et al., 2017), each component from the RL run was matched to a component from the LR run by identifying the maps with the highest degree of spatial correlation. We then analyzed the split-halves reliability using a Pearson correlation of the subject modes from the paired components.

Finally, we also assessed the reliability of the estimated independent components by repeating the process on the same data multiple times to ensure that our components do not represent a solution that is a local minima (Himberg et al.,2004). The Pearson correlation was analyzed for the spatial, temporal, and subject modes from corresponding components of different iterations.

## Results

The objective of our study was to find altered network responses associated in MDD compared to FHx and HC groups during tasks involving reward processing and perception of social stimuli. Our primary analyses utilized tensorial ICA (Beckman and Smith, 2005). For comparison, secondary analyses utilized conventional spatial ICA (McKeown et al., 1998) followed by dual regression (Filippini et al., 2009; Nickerson et al., 2017), both across the whole brain as well as restricted to striatum (Leech et al., 2012), given its importance in MDD (Shepard, 2013; Peters et al., 2016).

### Tensorial ICA reveals reward and social networks modulated by depression

Based on prior literature showing aberrant neural responses to reward in MDD (Ng et al., 2019) and social stimuli (Lai, 2014), we predicted that we would be able to identify differences associated with MDD in networks responsive to rewards and social stimuli. To test this hypothesis, we used tensorial ICA to capture systematic variation across participants (Beckman and Smith, 2005). We hypothesized that networks differentiating our HC, FHx, and MDD groups would contain clusters within the vmPFC, striatum, and cerebellum. For each task, the tensorial ICA was set to return 20 components each with a spatial, temporal, and subject mode representing the weight of each voxel, timepoint, and subject in the network. We manually classified components using common heuristics (Griffanti et al., 2017) and found that five components in the reward task and four components from the social task represented motion related noise. From the remaining components, we identified three networks that differentiated the MDD from HC group, all with spatial weights in the vmPFC, striatum, and cerebellum passing the gaussian mixture model threshold at a posterior probability of p>0.5. All three networks, one from the reward task and two from the social task, showed greater subject mode weights in HC compared to MDD indicating reduced network coherence in the MDD subjects: reward network one (Fig. 2) (*t*_(174)_ =-2.82 p=0.03 FWE d=-0.453 CI [-0.76, -0.14]), social network one (Fig. 3) (*t*_(174)_ =-2.78, p=0.04 FWE; *d*=-0. -0.43 CI [-0.72, -0.13]), and social network two (Fig. 4) (*t*_(174)_ =-2.75, p=0.04 FWE; *d*=- 0.429 CI [-0.73, -0.13]). However, there was no difference in engagement between HC from FHX groups: reward network one (*t*_(174)_ =-1.15, p=0.804 FWE), social network one (*t*_(174)_ =-0.66, p=0.99 FWE), and social network two (*t*_(174)_ =-.55, p=0.99 FWE). Nor did any of these networks identify differences between the MDD and FHx groups: reward network one (*t*_(174)_ =-1.78, p=0.4 FWE), social network one (*t*_(174)_ =-2.18, p=0.19 FWE), and social network two (*t*_(174)_ =-2.25, p=0.17 FWE). Additionally, none of the subject modes were correlated with motion as assessed by framewise displacement over the course of the task: reward network one (r=-0.03, p=0.63) social network one (r=-0.10, p=0.08), social network two (r=-0.04, p=0.48).

**Figure 2.**
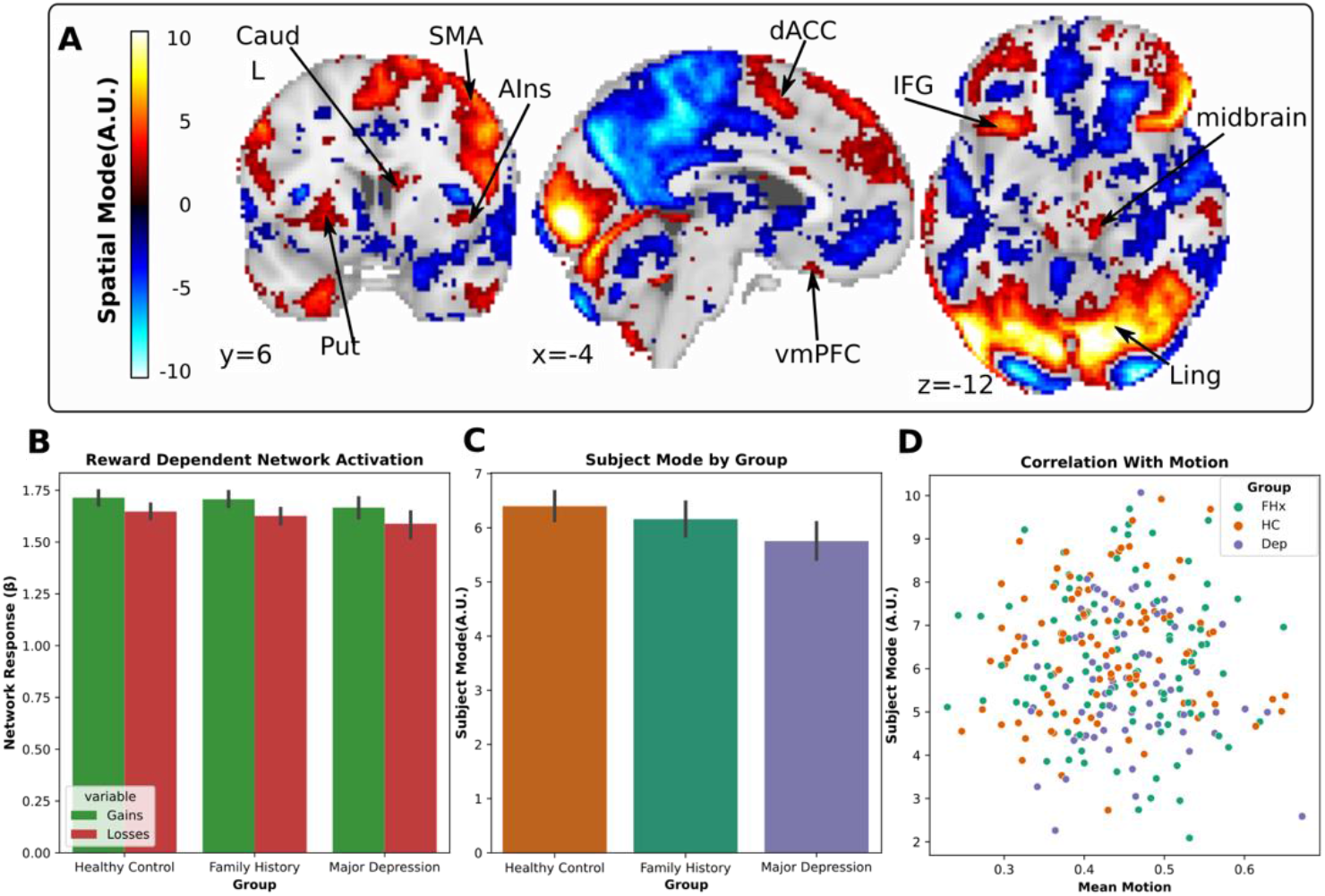
Depiction of reward network 08 revealed by tensorial ICA. (A) Spatial map of network weights with a threshold applied using gaussian mixture modeling highlights several regions that increase activation with the time course of the component including the vmPFC, dorsal anterior cingulate cortex (dACC), midbrain, inferior frontal gyrus (IFG), putamen (Put), and caudate (Caud). (B) The network showed greater activation in response to rewards than losses. (C) The greatest weight was associated with individuals from the HC group while individuals with MDD were the least weighted. (D) Individual weights with the subject mode were uncorrelated with motion.

**Figure 3.**
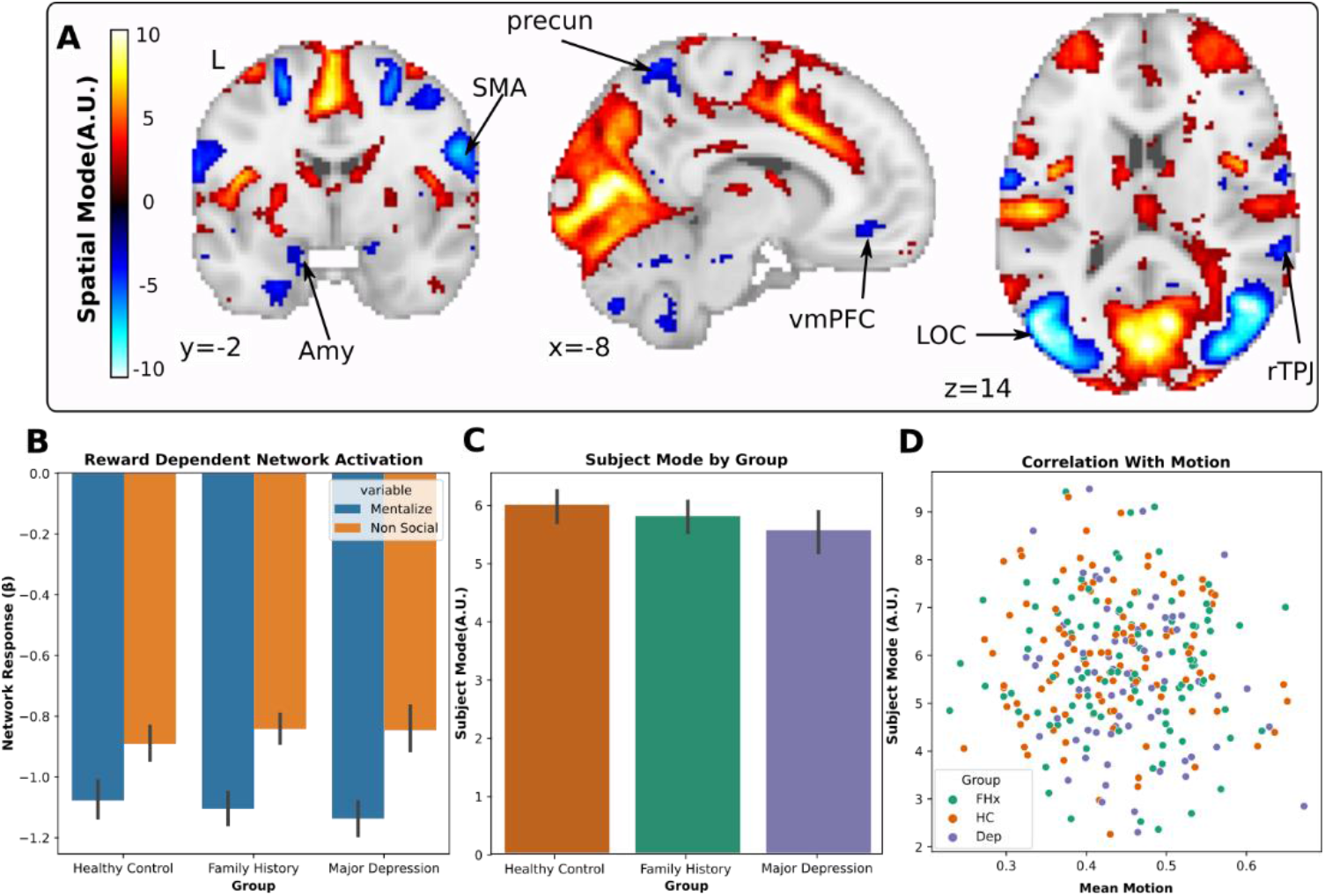
Depiction of social network 12 revealed by tensorial ICA. (A) Spatial map of network weights with a threshold applied using gaussian mixture modeling highlights several regions that decreased activation with the time course of the component including the vmPFC, precuneus (dACC), sensory motor area (SMA), right temporoparietal junction (rTPJ), amygdala (Amy), and lateral occipital cortex (LOC). (B) The network showed greater deactivation in response to mentalizing than non-social stimuli. (C) The greatest weight was associated with individuals from the HC group while individuals with MDD were the least weighted. (D) Individual weights with the subject mode were uncorrelated with motion.

**Figure 4.**
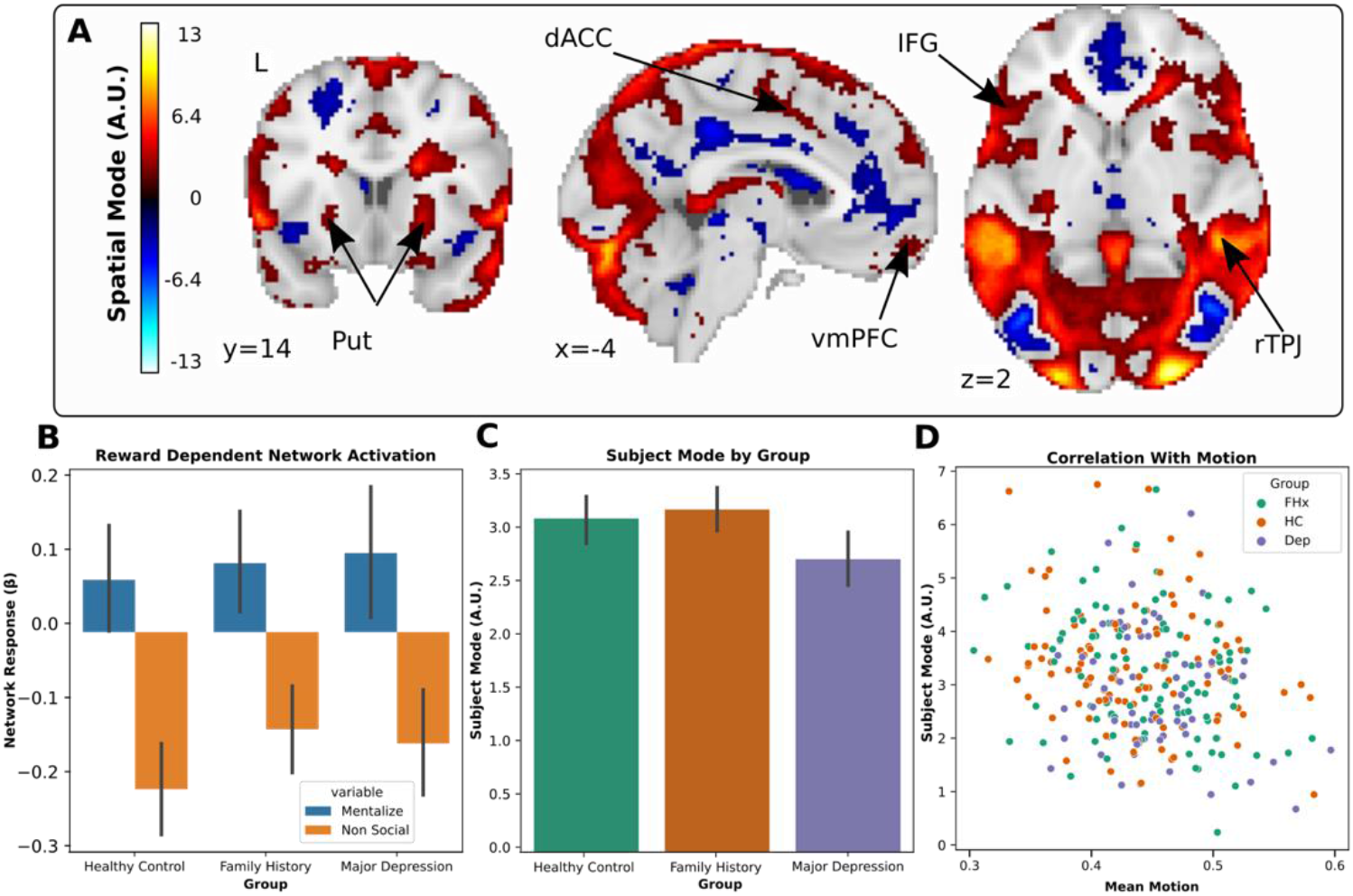
Depiction of social network 17 revealed by tensorial ICA. (A) Spatial map of network weights with a threshold applied using gaussian mixture modeling highlights several regions that increase activation with the time course of the component including the vmPFC, dorsal anterior cingulate cortex (dACC), inferior frontal gyrus (IFG), and putamen (Put). (B) The network showed activation in response to mentalizing condition and deactivation in response to the non-social condition. (C) The greatest weight was associated with individuals from the HC group while individuals with MDD were the least weighted. (D) Individual weights with the subject mode were uncorrelated with motion.

Next, we examined whether the group differences observed in network coherence were driven by responses to the task. To do this, we regressed the temporal mode of each component onto the blocks of different conditions in the gamble social cognition task and submitted the difference of the resulting betas to a series of two-sample t-tests. We found that the MDD compared to the HC group had a greater social-dependent activation of social network one (*t*_(174)_ =-2.78 p=0.049 FWE d=-0.43 CI [- 0.74, -0.12]) which included clusters in the vmPFC, precuneus, cerebellum, and amygdala (p>0.5 GMM) which increased in response to social stimuli. Additionally, exploratory analyses showed that all three networks had strong task-dependent activation across all participants: reward network one (Fig 2.) (*t*_(174)_ =6.07, p<0.0001 FWE; *d*=-0.514 CI [0.35, 0.68]), social network one (Fig 3) (*t*_(174)_ =- 16.5, p<0.0001 FWE; *d*=-1.38 CI[-1.55,-1.16]), and social network two (Fig 4) (*t*_(174)_ =16.61, p<0.0001 FWE; *d*=-1.41 CI[1.14,1.62]). This suggests that while all three networks were related to the task, the differences between groups in social network one were driven, at least in part, due to differences in task-dependent network activation.

We additionally took steps to analyze the possible reliability and reproducibility of our results. First, we gauged the measurement reliability of our tensor ICA components by conducting a test to measure the split-halves reliability. The mean correlation of subject modes between both runs for components identified from the Social task was r=0.59 and the mean for components from the Gamble task was r=0.63. These results indicate that the tensorial ICA results derived from the HCP data are reproducible across runs.

We also performed additional measures of validation were performed to assess the plausible reproducibility of our results in other samples. Using a linear discriminant analysis with 3-fold cross validation and permutation analysis to identify p-values if we were able to discriminate MDD from HC participants. The subject mode of the identified reward network had 65% accuracy(p<0.001) while the first social network showed 57% accuracy (p=0.92) and second social network 62% accuracy (p=0.012). This suggests that two of the three components we identified should be reproducible out of our limited sample.

Finally, we correlated iterations of the tensorial-ICA process on the same data to ensure that our components represented optimal solutions for the ICA process. Corresponding components from both Social and Gamble task data were highly correlated in the spatial, temporal and subject modes (all r>0.99). These results indicate highly stable components within the ICA framework.

### Whole brain dual regression reveals no networks modulated by depression

We followed our results of the tensorial ICA with a whole-brain dual regression (Filippini et al., 2009; Nickerson et al., 2017) to better contextualize and assess the effectiveness of the tensorial ICA networks in identifying individual differences associated with MDD. The initial ICA from the dual regression was set to return 25 components, but we chose to reduce comparisons by selecting only 10 of these component networks. We chose these 10 networks by identifying the components that showed the greatest spatial correlations to 10 canonical brain networks identified in prior work (see Table 1) (Smith, 2009). These networks have also been linked to alterations in resting-state connectivity in patients with depression (Mulders et al., 2015; Cieri et al., 2017).

First, we tested whether the activation of the networks revealed by the whole-brain dual regression were modulated by task by regressing the time-course of each network onto the gains and losses and submitted the difference of the resulting beta weights to a one sample t-test. We found that three of our canonical brain networks showed a reward-dependent activation, including the networks most closely resembling the Vis1 (*t*_(278)_ = 5.938, p< 0.0001), Vis3 (*t*_(278)_ = 10.013, p < 0.0001), and left frontoparietal network (*t*_(278)_ = 6.682, p< 0.0001) (Fig. 5). However, all 10 canonical brain networks showed differential activation in response to social stimuli (all p <0.0014 FWE) (Fig. 6). Next, to determine if any of these networks were modulated by clinical status, we submitted the difference between the task beta weights to a two-sample t-test. We found that differences between our MDD, FHx, and HC groups in reward or social activation across any of the 10 canonical (all p>0.05). Since no effects were shown across our ten canonical networks, we expanded our analysis of between group effects to include all 25 components for both social and reward tasks. We found no significant effects between our MDD, FHx, and HC groups in reward or social activation across any of our component networks (all p>0.05).

**Figure 5.**
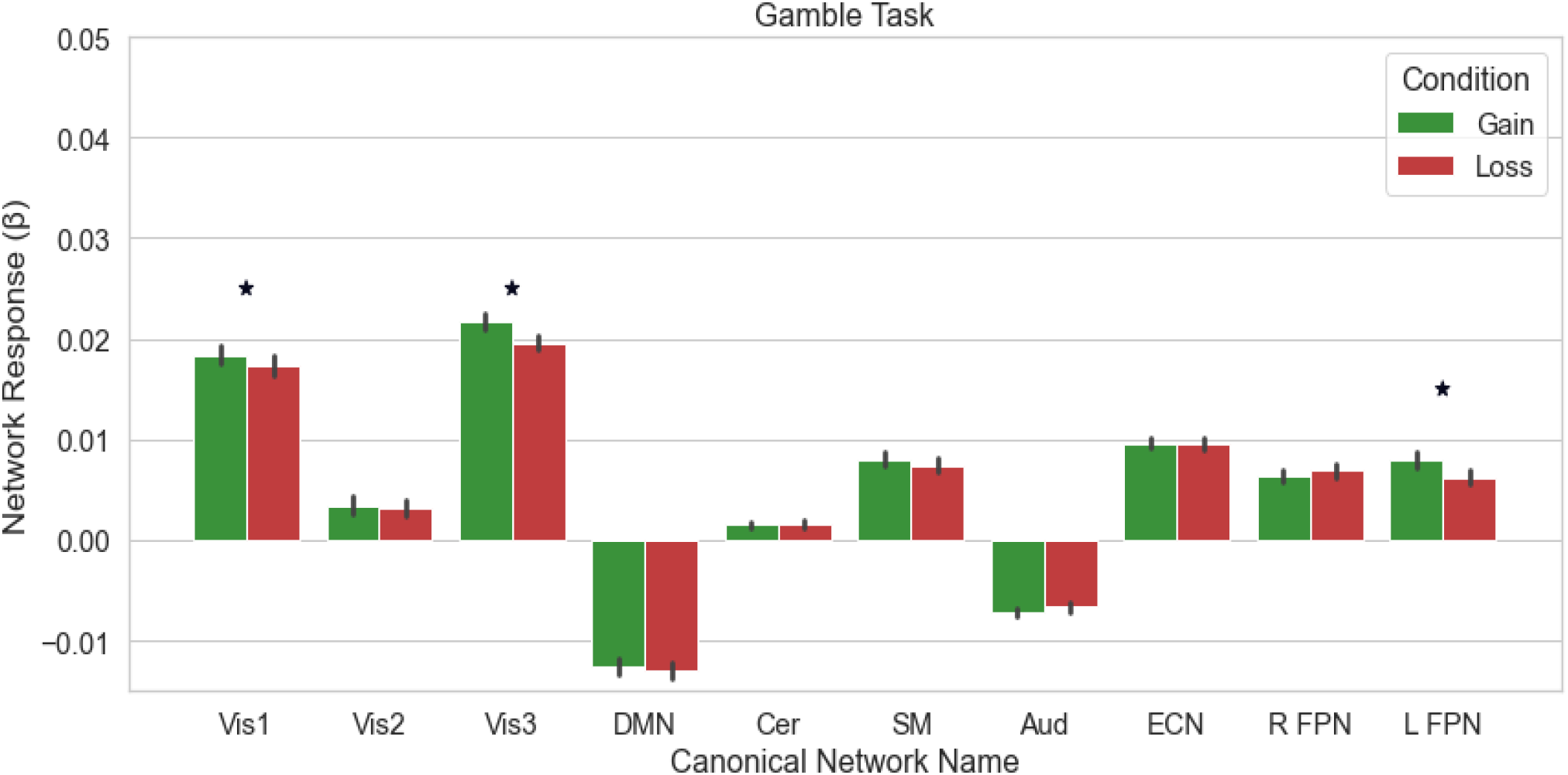
Dual regression revealed three canonical component networks were modified by reward. The three networks whose spatial maps was most closely correlated with the Vis1 (r=0.76), Vis 3 (r=0.59), and LFPN (r=0.65) canonical networks were significantly different activation during reward and loss blocks.

**Figure 6.**
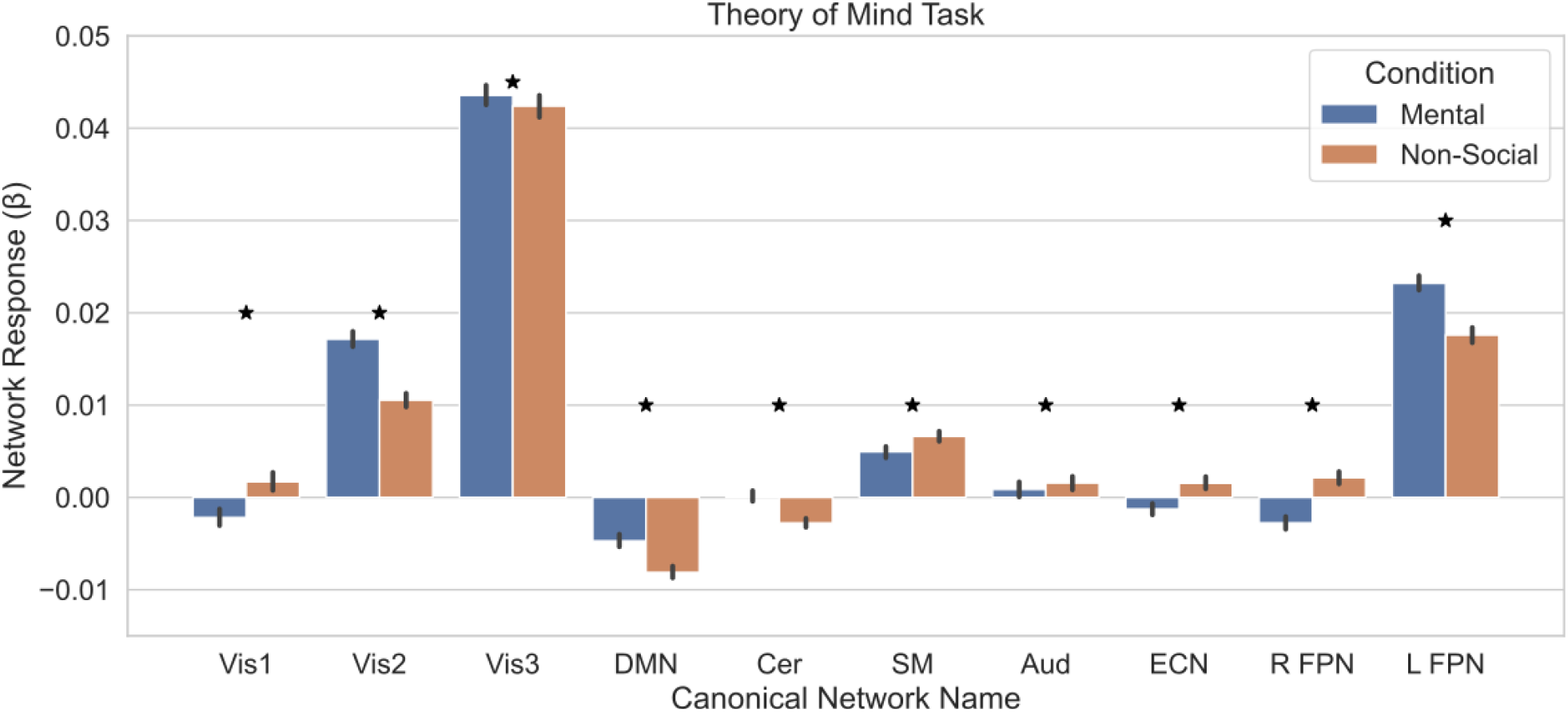
Dual regression revealed all 10 canonical component networks were modified by social stimuli. The responses to the mental and non-social conditions are shown for each of the component networks which were most highly correlated with our canonical brain networks.

### Spatially restricted dual regression reveals enhanced social network activation in depression

Analyses from the whole-brain dual regression revealed no networks with task-dependent activation modulated by depression. However, other modifications to ICA might still be able to find some networks with activation modulated by depression without taking advantage of the between subject variance offered by tensorial ICA. For example, previous work has used dual regression following ICA on data masked to the posterior cingulate cortex (PCC) to find networks mapped to the complex architecture of the PCC (Leech et al., 2012). We used a similar method, using dual regression following ICA on data masked to the striatum, to take advantage of the weakened cortico-striatal structural connections associated with depression (Bracht et al., 2015). Additionally, the involvement of cortico-striatal networks in reward processing (Smith et al., 2016) and social behavior (Molesworth et al., 2015; Bentea et al., 2021) suggested that we could take advantage of these effects to identify task-dependent networks. We predicted that by spatially restricting the ICA step of our dual regression (Leech et al., 2012) to the striatum, we would be able to identify networks with task-dependent activation modulated by MDD.

The initial ICA was restricted to the striatum and set to return 25 components. After spatial regression identified individual time courses of each striatal component, the temporal regression across the whole brain revealed whole brain networks showing connectivity with each component of the striatum. After averaging these individual whole-brain maps, we took the same approach as the whole-brain dual regression to identify ten canonical brain networks (Table 1). Though we had expected each canonical network to be represented by unique components from the spatially restricted dual regression, it is worth noting that two pairs of networks were represented by the same component. In the reward task, one component represented both the Vis1 and Vis3 networks and another component represented both the DMN and Aud networks. In the social task, one component represented both the Vis2 and right FPN (associated with guided attention) (Fisher et al., 2020) networks and another component represented both the DMN and ECN networks. This reduced the number of comparisons made from ten to eight in our analyses across both tasks. Additionally, individuals should be careful to not interpret claims about these specific components as claims about each or both networks separately.

To assess the reward-dependent activation of the networks from the spatially restricted dual regression we regressed the time-course of each network, for each task, onto the condition block design and submitted the difference of the resulting beta weights to a one sample t-test. We found that none of the networks were responsive in the reward task (Fig. 7) (all p > 0.05 FWE), but 3 networks were responsive in the social stimuli, including the components most closely correlated with the vis1 (*t*_(278)_ = 7.49 p<0.0001) network, left FPN network (*t*_(278)_ = -4.26, p<0.0001), and DMN/ECN (*t*_(278)_ = 4.22, p<0.0001) network component. We found none of the 10 canonical networks from the reward task (all p>0.05 FWE) or the social task (all p>0.05) revealed an interaction between task activation and clinical status. As none of our canonical brain networks revealed an interaction between task activation and clinical status, we expanded our analyses to include the full set of 25 component networks from both tasks.

**Figure 7.**
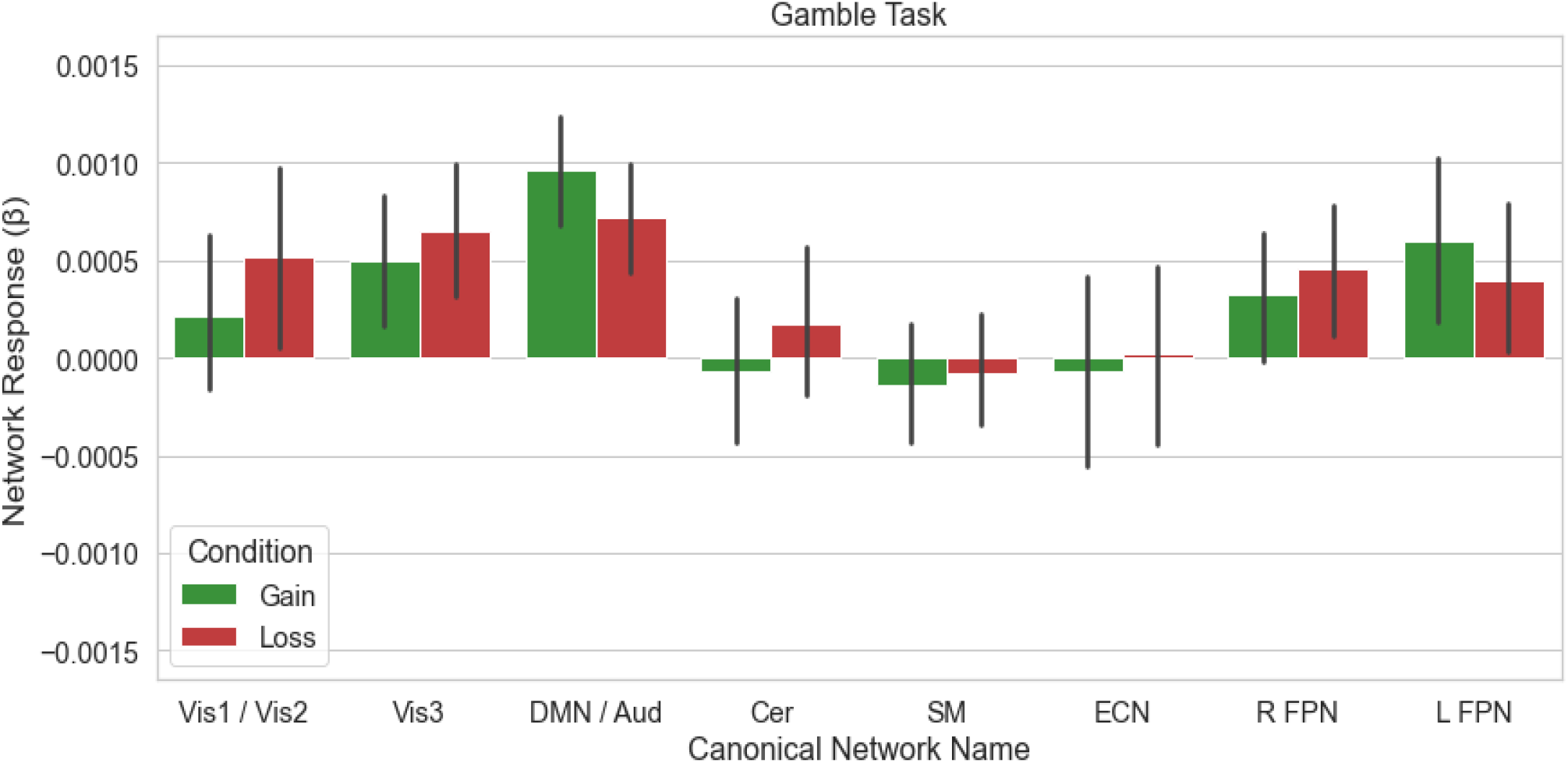
Spatially-restricted dual regression revealed no component networks that were modified by rewards. The responses to the gain and loss conditions are shown for each of the component networks which were most highly correlated with our canonical networks.

Expanding our analyses revealed a cortico-striatal network, represented by social component 09 (Fig. 9.), with a coactivation between subregions of the striatum and vmPFC that had a significantly greater task dependent activation in the MDD than in the HC group (*t*_(174)_ =3.459 p=0.0177 FWE). In order to better understand the task-dependent activation of our network across each group we conducted three post-hoc tests. Post hoc comparisons revealed that while network activation was greater in the non-social than mental condition for participants in the MDD group (*t*_(70)_ = 3.691 p=0.0004), there was no significant task-dependent activation for the HC (*t*_(102)_ = 0.96 p=0.337) and FHx group (*t*_(104)_ =1.598 p=0.113).

**Figure 8.**
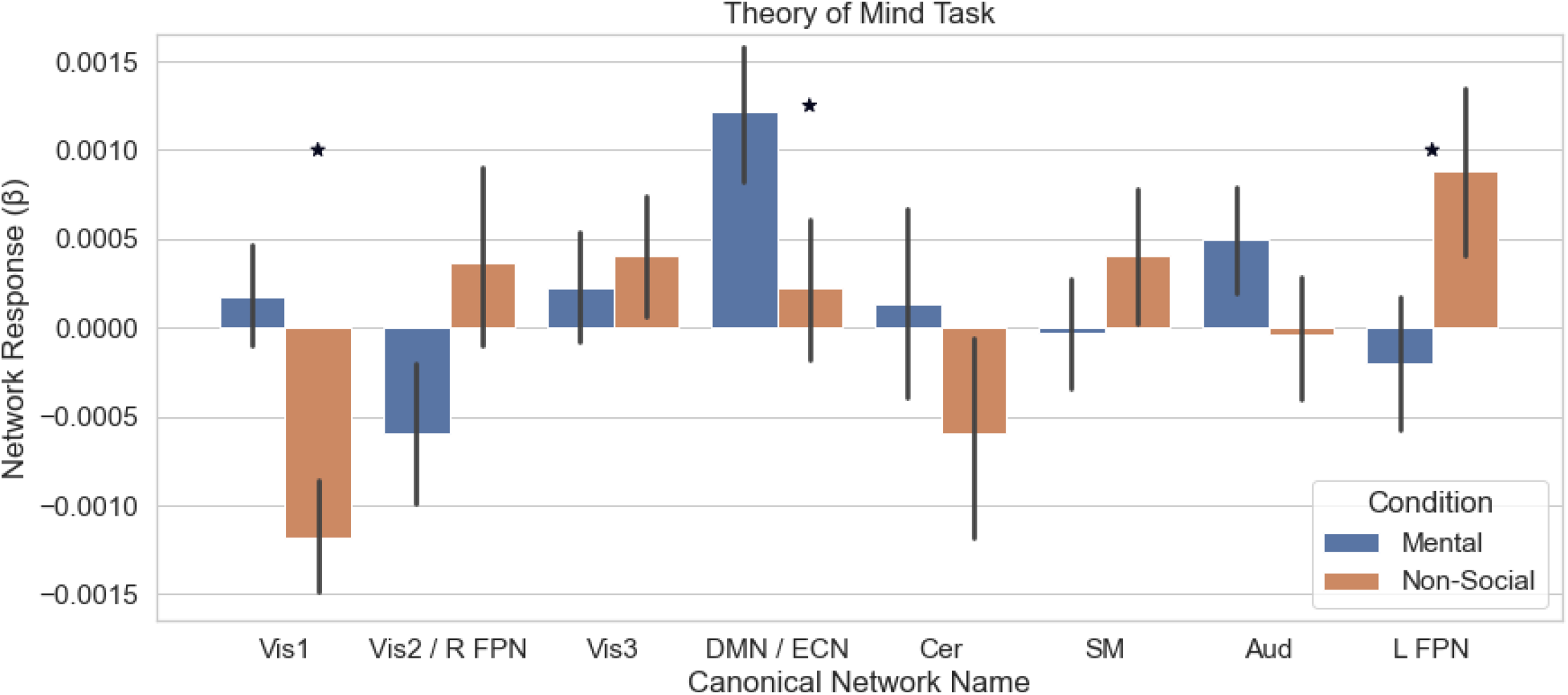
Spatially-restricted dual regression revealed three canonical component networks modified by social stimuli. The responses to the mental and non-social conditions are shown for each of the canonical component networks. A significant difference in activation between conditions was observed for the Vis1, DMN/ECN, and L FPN networks.

**Figure 9.**
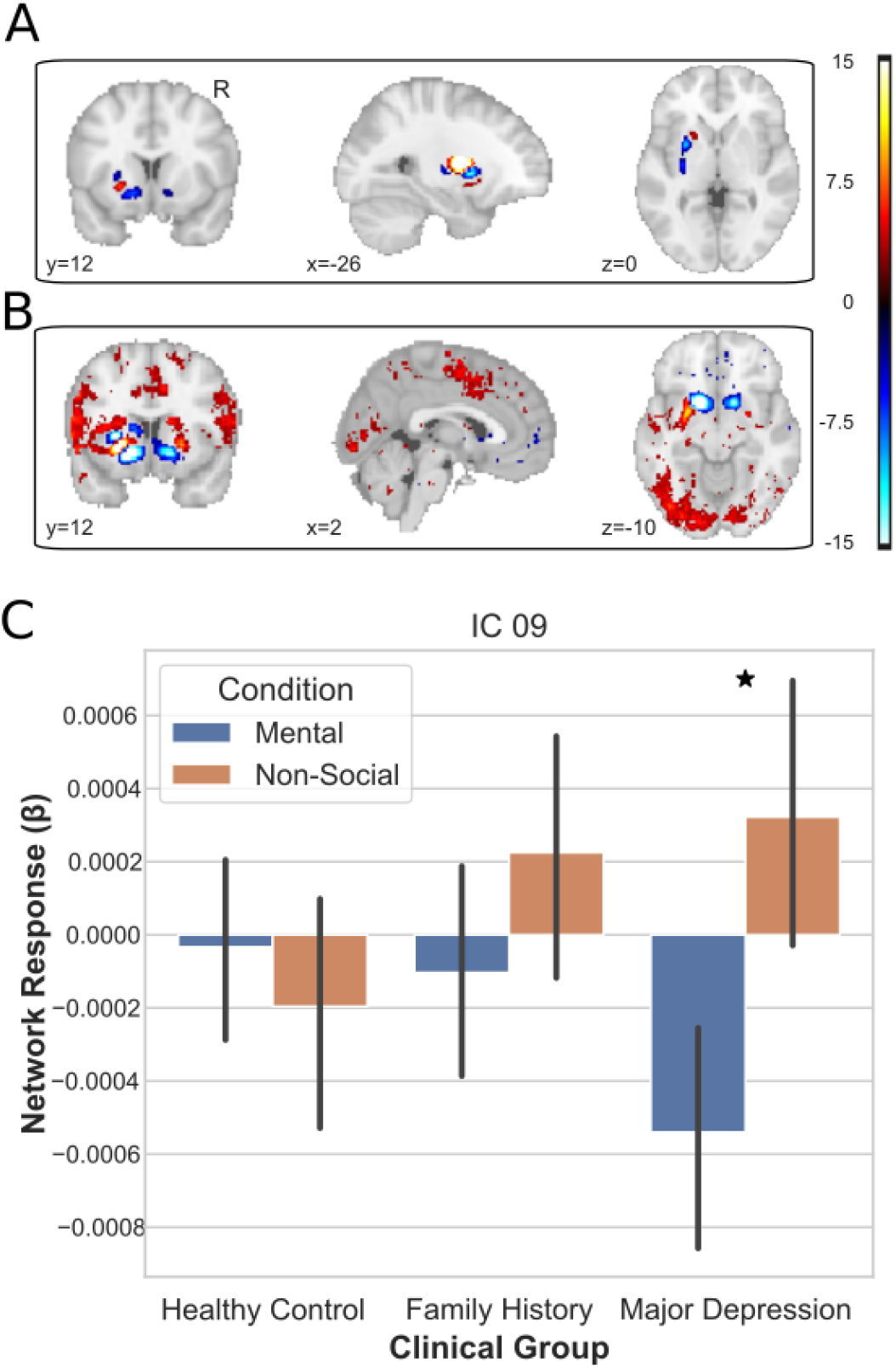
Depiction of clinically sensitive social network from the spatially restricted dual regression. (A) The projection of weights from the spatially restricted ICA onto the striatum. (B) Results of temporal regression reveals connections between cortex and spatial mode of the striatum. (C) The MDD group was the only group to show differential activation of component 09 in response to the social and non-social stimuli.

## Discussion

Functional connectivity has been commonly used to identify neural differences associated with psychiatric disorders (Buckner et al., 2013). Common models of functional connectivity, using spatial or temporal ICA in the initial steps ignore sources of inter-subject variability that may be crucial for identifying individual differences. The objective of our study was to examine whether tensorial ICA, by including sources of inter-subject variability, could identify corticostriatal networks that differentiate participants with a prior diagnosis of depression, a family history of depression, and healthy controls when applied to functional data from a gambling and social cognition task. To meet our objective, we analyzed data from the Human Connectome Project using tensorial ICA and compared our results to those obtained using a whole-brain and spatially restricted dual regression. Using tensorial ICA we found 3 different corticostriatal network maps that differentiated our MDD sample from healthy controls. This effect was in contrast to the whole brain dual regression which revealed no group differences and the spatially restricted dual regression which revealed one social network that showed differential activation between the MDD population and controls.

Our results identified 3 networks with decreased coherence in MDD, each revealing connections to regions across the brain similar to both the default mode network (DMN) (medial PFC, PCC, and angular gyrus) and salience network (insula, thalamus, striatum) consistent with prior work that targeted MDD associated changes in connectivity in the DMN (Høgestøl et al., 2019; Zhang et al., 2016) and salience networks (Godfrey et al., 2022) as key factors in MDD. We also showed that lower coherence in a reward network containing striatum, vmPFC, and anterior cingulate cortex was associated with MDD, consistent with seed-based results showing that MDD modulated striatum-vmPFC (Liu et al., 2021; Hanson et al., 2018) and striatum-cingulate connectivity (Admon et al., 2015; Walsh et al., 2017) in response to reward. The social networks we found with lower coherence in MDD also contained striatum, vmPFC, and cingulate cortex, consistent with suggestions that MDD altered connectivity during social processing between the striatum and vmPFC (Healy et al., 2014; Tepfer et al., 2019) as well as between the striatum and cingulate (Wang et al., 2015). Inconsistent with results of MDD connectivity in social processing however, neither of the social networks we identified contained the Nacc subregion of the striatum as suggested in prior work (Healy et al, 2014; Wang et al., 2015; Tepfer et al., 2021). Additionally, these previous results used methods that were only able to identify one network while our results suggest that there are disruptions in two independent social networks associated with MDD.

The functional connectivity associated with major depression has been studied using several methods, including dual-regression and other analyses relying on ICA (Taylor et al., 2021). However, traditional methods of using ICA often ignore between subject variability. Our results suggest that including the between subject variability using tensor ICA could be used to better identify network responses that are associated with MDD. One previous study had successfully applied tensor ICA to identify functional connectivity differences associated with dementia and concluded that it provided a significant benefit over traditional general linear model approaches. Our study was both the first to apply tensorial ICA to identify functional network differences associated in major depressive disorder and to compare those results to another ICA based method (dual regression). Despite the previous success applying tensor ICA to dementia, which often presents in a similar neurological profile of frontotemporal degeneration (Bang et al., 2015), no one had tested whether tensor ICA would provide similar benefits for MDD, which has a more heterogenous neurological profile (Dinga et al., 2019; Drysdale et al., 2017). Additionally, it was not certain that there would be a benefit over traditional ICA, as both take a multivariate approach that has been shown to be especially sensitive to task and group differences (Norman et al., 2006; Haxby, 2012)

While this study presents a case for the use of tensorial ICA in detecting individual differences in functional connectivity, there are several limitations to our study. The first limitation is drawn from the existence of numerous subtypes of depression (Drysdale et al., 2017) and the heterogeneity of experiences in depression. We had limited access to the clinical information available and only attempted to analyze differences associated with lifetime diagnosis of depression or a family member with a lifetime diagnosis. Our results do not speak to the multiple possible combinations of depression symptoms or account for whether a subject was experiencing a depressive episode at the time of their scan. However, the Human Connectome Project has recently started new protocols including one specifically aimed at understanding the cross-cutting psychiatric features of depression and anxiety disorders (Tozzi et al., 2020). This new data from the HCP may help to analyze subtypes and other features of depression. Second, while we chose two tasks that were associated with deficits in depression, the lack of robust behavioral measures and block design of these tasks poses significant limitations on interpretation for behavior. While our results showed that one reward and two social networks were modulated by depression, we are unable to speak to what processes these networks might reflect. Ideally, we would like to show that the coherence or activation of our tensor networks mediated the effect of depression on a specific behavior associated with MDD. Showing an effect on behavior in the reward task is not feasible as there is no interpretable behavior in the task (only random guesses). In the social task, previous work relying on the same sample showed that MDD participants marked a larger number of videos as social (Tepfer et al., 2020). However, the block design of this task makes it difficult to analyze how the network activity is tied to specific trials (Josephs & Henson, 1999) which is necessary to show the relationship between network activity and behavior on specific trials.

Additional limitations are related to difficulties of reproducing brain related markers of clinical disorders (Woo et al., 2017). Even within a large open data set such as the human connectome project we were unable to identify a large enough sample of individuals with MDD to perform an independent validation of our results. Future studies could use the spatial maps identified in our study (available on NeuroVault: https://identifiers.org/neurovault.collection:13474) and assess how well they distinguish MDD and healthy controls in independent datasets involving reward processing and/or social cognition. Especially, as small samples are at risk of overestimating differences between groups, these results should be taken to represent an initial description of the networks involved in depression and not a demonstration of diagnostic capability, which would require larger samples and prospective validation (Woo et al., 2017). Finally, we note that future studies should assess the test-retest reliability of tensorial ICA in the context of individual differences in psychopathology (Zuo et al., 2010). Quantifying this form of reliability for tensorial ICA would be an important step forward clinical studies seeking to leverage this tool to study individual differences in task-dependent brain networks.

We were able to use tensorial ICA to identify activation of three cortico-striatal networks altered by depression. Although each of these networks were statistically independent, they shared overlapping representation within the vmPFC, striatum, cerebellum, and cingulate cortex. This suggests that the striatum, cerebellum, vmPFC, and cingulate cortex may form a network that is a common feature of aberrant reward and social processing in MDD. Additionally, given the structural connections of the striatum to the cerebellum (Bostan & Strick, 2018) as well as the cingulate and vmPFC (Haber and Knutson, 2010) it seems as though the striatum may play a role as the central hub in this network.

Additionally, using tensorial ICA provided a significant advantage over traditional ICA which revealed fewer differences, suggesting there should be effort to work past the limitations of a common time course among subjects and take advantage of the technique. For studies that already meet this common time course requirement, tensor ICA could be used for potentially better denoising solutions or feature selection prior to multivariate pattern analysis by taking the place of temporal or spatial ICA. Many advances have been made by exploring the functional connectivity related to psychiatric disorders (Finn and Constable, 2022; Woodward and Cascio, 2015) and tensorial ICA has the potential to be a powerful tool to continue these advances.

## Acknowledgments

This work was supported, in part, by grants from the National Institutes of Health (R03-DA046733 and RF1-AG067011 to DVS). We thank Vishnu Murty and Thomas Olino for offering constructive suggestions during the preparation of this manuscript. Data collection and sharing for this project was provided by the Human Connectome Project (HCP; Principal Investigators: Bruce Rosen, M.D., Ph.D., Arthur W. Toga, Ph.D., Van J. Weeden, MD). HCP funding was provided by the National Institute of Dental and Craniofacial Research (NIDCR), the National Institute of Mental Health (NIMH), and the National Institute of Neurological Disorders and Stroke (NINDS). HCP data are disseminated by the Laboratory of Neuro Imaging at the University of Southern California.

